# BENEFICIAL EFFECTS OF KETOGENIC DIETS FOR CANCER PATIENTS – A REALIST REVIEW WITH FOCUS ON EVIDENCE AND CONFIRMATION

**DOI:** 10.1101/137950

**Authors:** Rainer J. Klement

## Abstract

**Background:** Ketogenic diets (KDs) have gained popularity among patients and researchers alike due to their putative anti-tumor mechanisms. However, the question remains which conclusions can be drawn from the available human data thus far concerning the safety and efficacy of KDs for cancer patients.

**Methods:** A realist review utilizing a matrix-analytical approach was conducted according the RAMEsEs publication standards. All available human studies were systematically analyzed and supplemented with results from animal studies. Evidence and confirmation were treated as separate concepts.

**Results:** 29 animal and 24 human studies were included in the analysis. The majority of animal studies (72%) yielded evidence for an anti-tumor effect of KDs. Evidential support for such effects in humans was weak and limited to individual cases, but a probabilistic argument shows that the available data strengthen the belief in the anti-tumor effect hypothesis at least for some individuals. Evidence for pro-tumor effects was lacking completely.

**Conclusions:** Feasibility of KDs for cancer patients has been shown in various contexts. The probability of achieving an anti-tumor effect seems greater than that of causing serious side effects when offering KDs to cancer patients. Future controlled trials would provide stronger evidence for or against the anti-tumor effect hypothesis.

## 1. Introduction

Ketogenic diets (KDs) are diets that mimic the metabolic state of fasting by inducing a physiological rise in the two main circulating ketone bodies, acetoacetate and beta-hydroxybutyrate (BHB), above the reference range (typically ≥ 0.5 mmol/l for BHB). KDs have recently gained increased interest for the treatment of a variety of diseases, where they are used either as a stand-alone metabolic therapy or as part of a broader therapeutic approach [1]. One of these diseases is cancer, a chronic systemic disease characterized by dysfunctional mitochondria and an increased dependence on substrate fermentation (in particular glucose) as against oxidative phosphorylation. Theoretically this provides an ideal target for ketogenic metabolic therapies [2–4]. While mechanistic reasoning based on experimental work and preliminary human studies provide some support for KDs in cancer treatment [5–9], it has been criticized that there is no evidence for KDs being beneficial for cancer patients due to the lack of randomized controlled trials (RCTs). For example, Huebner et al. [10] stated: *“Following scientific practice e.g. with herbs, controlledif possible randomized clinical studies are mandatory as scientific evidence for the efficacy of a diet and the basis of a positive recommendation. Risks may be derived from case reports and even preclinical data.”* However, such statements appear problematic for at least four reasons. First, it draws a bad analogy between interventions with single agents such as herbs and a complex dietary change. Second, it does not allow for integrating patient values or patient circumstances [11] when counseling individual patients. Third, it neglects the possibility of high-quality mechanistic reasoning as another potential source of evidence [11]. Fourth, and more generally, this hierarchical view of evidence, the hallmark of the evidence based medicine paradigm, is problematic due to the conceptual incompatibility of external and internal validity which cannot both be maximized simultaneously [12,13]. It should be considered that different research methods complement each other because the types of validity they produce are incompatible and complementary [13]. The reader is invited to do her own thought experiment about how a RCT to test the effects of a KD for cancer patients should be conducted. How should the inclusion and exclusion criteria be formulated? More generally, would findings concerning patients who fulfilled the inclusion criteria be translatable to patients who were or would have been excluded from the study? A recent clinical study testing a KD in advanced cancer patients exemplifies this problem. In this study by Tan-Shalaby et al. [14], most patients had to be recruited from locations more than a 3h drive away: *“Majority of the interested patients who lived locally did not qualify, as they were not US veterans, one reason for inclusion.”* [14] Could such a long way to the hospital not possibly confound the results, e.g. due to increased cortisol levels associated with the stress of traveling? This example reveals one of several problems of placing RCTs over other types of studies for deriving evidence, relating to internal versus external validity, specific versus overall treatment effects, and context dependence of the intervention [12]. Several studies on the KD in cancer patients have already been published; to simply dismiss all those as “no evidence” makes the life of the investigator easy, but is epistemologically invalid because it confuses the concepts of data and evidence. Studies by themselves do not constitute evidence but deliver data which are context free; it is the data that *can* constitute evidence but only *within a context provided by competing hypothesis* [15]. Therefore, even small studies and case reports can constitute evidence (although with *a priori* low strength [11]) if they are relevant for discriminating between two competing hypotheses. Neglecting them would violate the principle of total evidence [16]^1^.

In an alternative approach, one could therefore collect all available data in order to try answering the questions “does a KD work, and if yes, for which cancer patients, under what circumstances, and how?” Such an approach is naturally pursued within the philosophy of realism [17–19], which states that *“there is a world that exists independently of our perception of it, and that our theories inform us about the existence and nature of this realm”* [20]. A realist review emphasizes inquiry about the relationship of an intervention with the outcome dependent on the mechanisms that connect them and the context in which this relationship occurs [17].

This work therefore aims to present a systematic realist synthesis of the question whether KDs are beneficial for cancer patients. The review will be complemented by a matrix analysis in order to collate the totality of available evidence [13]. The following review questions were formulated to guide the analysis: (1) Given the data, to what degree should we believe that KDs exert anti-tumor effects in cancer patients? (2) What is the evidence for the hypothesis that KDs have beneficial effects for cancer patients both in terms of an anti-tumor effect as well as improving quality of life (QoL)? (3) What are the mechanisms that researchers put forward to explain their findings? We term the first question the *confirmation* question, the second the *evidence* question [15] and the third the *causation* question.

This review was conducted and structured according to the RAMESES reporting standards for realist synthesis [19].

## 2. Materials and Methods

Realist reviews were originally designed, and still are mainly used for investigating complex policy interventions. However, a KD intervention is certainly also too complex to be compared with a pharmaceutical intervention such as a targeted therapy since it requires behavioral changes in eating patterns, induces complex metabolic changes impacting a multitude of cell signaling pathways and can be implemented in a great variety of ways (examples are the Spanish Mediterranean KD [21], Paleolithic KD [22], Medium chain triglyceride KD [23] or even a vegan version [24]). I therefore chose a realist review approach, but adapted towards a more qualitative analysis by collating all individual research findings in a matrix form. This combination of realist synthesis with matrix analysis was proposed by Walach and Loef [13] and is especially useful for the evaluation of lifestyle and alternative and complementary medicine interventions.

It was planned to utilize every study in which cancer patients had been treated with a KD based on an anti-tumor rationale. Additionally, animal studies were planned to be investigated in order to gain insights into mechanistic reasoning; thereby only those studies in which the diet was started on the day of or after tumor implantation were considered since these mimic the clinical situation in which a KD is introduced after cancer is already manifested. The primary search was conducted in Pubmed on May 10^th^ 2017 using the term “ketogenic diet AND cancer". A secondary search was conducted in Scopus, reference lists and a personal electronic library (https://www.mendeley.com). From the animal studies the tumor model, concurrent treatment modality and outcome was recorded. From the studies on humans, I extracted the study type, number of patients treated, tumor location and stage, type of KD (calorie restricted or not), concurrent treatment, as well as outcomes, context and proposed mechanisms as applying to the three research questions formulated above.

For answering these questions, a distinction must be made between evidence and confirmation as pointed out by Bandyopadhyay et al. [15]. While confirmation measures the degree of belief in a single hypothesis, the evidence is always evaluated against a second hypothesis. Given some background knowledge or context C, the data D are interpreted as *evidence* for a hypothesis H_1_ as against hypothesis H_2_ if the likelihood ratio P(D|H_1_ & C)/P(D|H_2_ & C)>1; in contrast D *confirm* a hypothesis H if and only if they raise the posterior probability of H compared to its prior probability, P(H|D)>P(H) [15]. In this way, H_1_is better confirmed by the data than H_2_if P(H_1_|D) > P(H_2_|D). From Bayes theorem, P(H|D)=P(H)*P(D|H)/P(D), this also implies that in case of a high prior probability P(H) for a hypothesis H, we would stick to believing in H even if there only are few observations supporting it, while in case of a low P(H) observational support for H would have to be large in order to convince us to believe in it [25]. A prior probability for H could be derived, e.g. from preclinical studies [13]. To answer the evidence question, which implies contrasting two hypotheses against each other, the anti-tumor effect hypothesis (predicting effects against tumor growth, metabolism or genetics) and the pro-tumor effect hypothesis (predicting tumor growth promotion) were each compared to the hypothesis of a neutral or no effect, respectively. Similarly, the positive QoL effect hypothesis (hypothesizing an improvement of physical, psychological or emotional functioning through a KD) and the negative QoL effect hypothesis (predicting a QoL detriment) were each compared to the hypothesis that a KD does not affect QoL. Given the non-analytical form of these hypotheses, the judgment of the evidence remained qualitative in the sense that I only asked whether - *ceteris paribus* - the individual study data appear more likely under the considered hypothesis as against its competitor hypothesis or not. The methods, results and discussion sections of the individual studies were used to guide this decision under the consideration of the specific context.

## 3. Results

The primary search in PubMed resulted in 157 articles of which 13 where original research papers on tumor patients [14,26–37], 32 on animals [38–69], two studies on both [70,71], and two were meta-analysis of animal studies [72,73]. Another seven animal studies [74–80] and nine human studies [81–89] were identified by the secondary search process. Subsequently, 14 animal studies were excluded because they either initiated the KD prior to tumor cell injection [43,45,68,69], had no control group [46] or because they basically replicated the results of a previously published tumor model system ([39,40,75] replicated [38], [76] replicated [44], [59] replicated [77], [61] replicated [53], [80] replicated [58], and [62, 67] replicated [47,49]). A flow chart of the search process is given in Figure 1.

**Figure 1.**
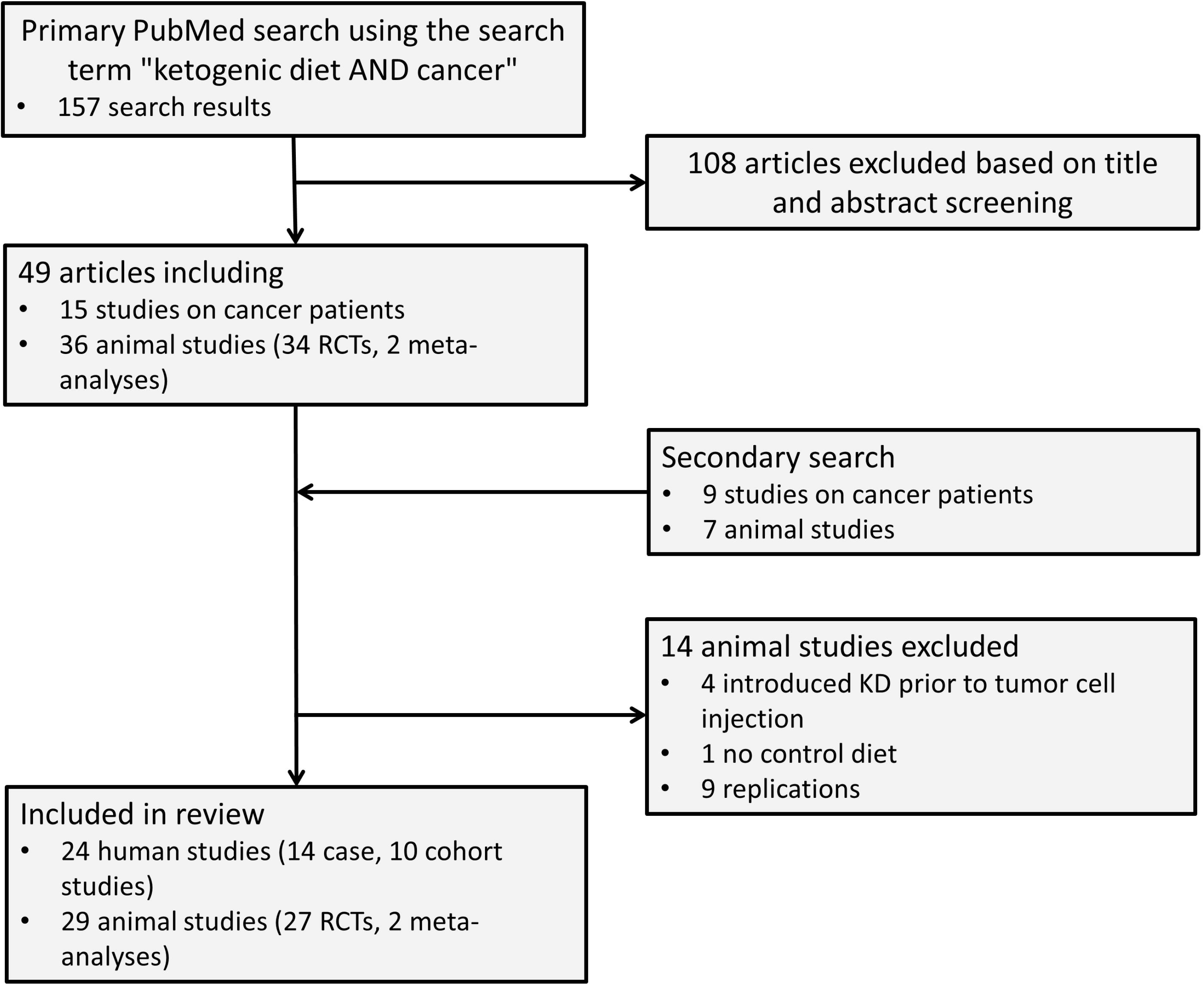
Flow chart of the study acquisition process.

Table 1 provides an overview of the selected patient studies with a focus on the efficacy and QoL outcomes; further details are given in Supplementary Table 1. There were 14 case studies and 10 cohort studies in which a total of 214 patients had been treated with a KD. The results concerning evidence for and against anti-tumor effects of the KD are tabulated in Table 2 together with the results of the animal studies. Details of these animal experiments are given in Supplementary Table 2.

**Table 1.**
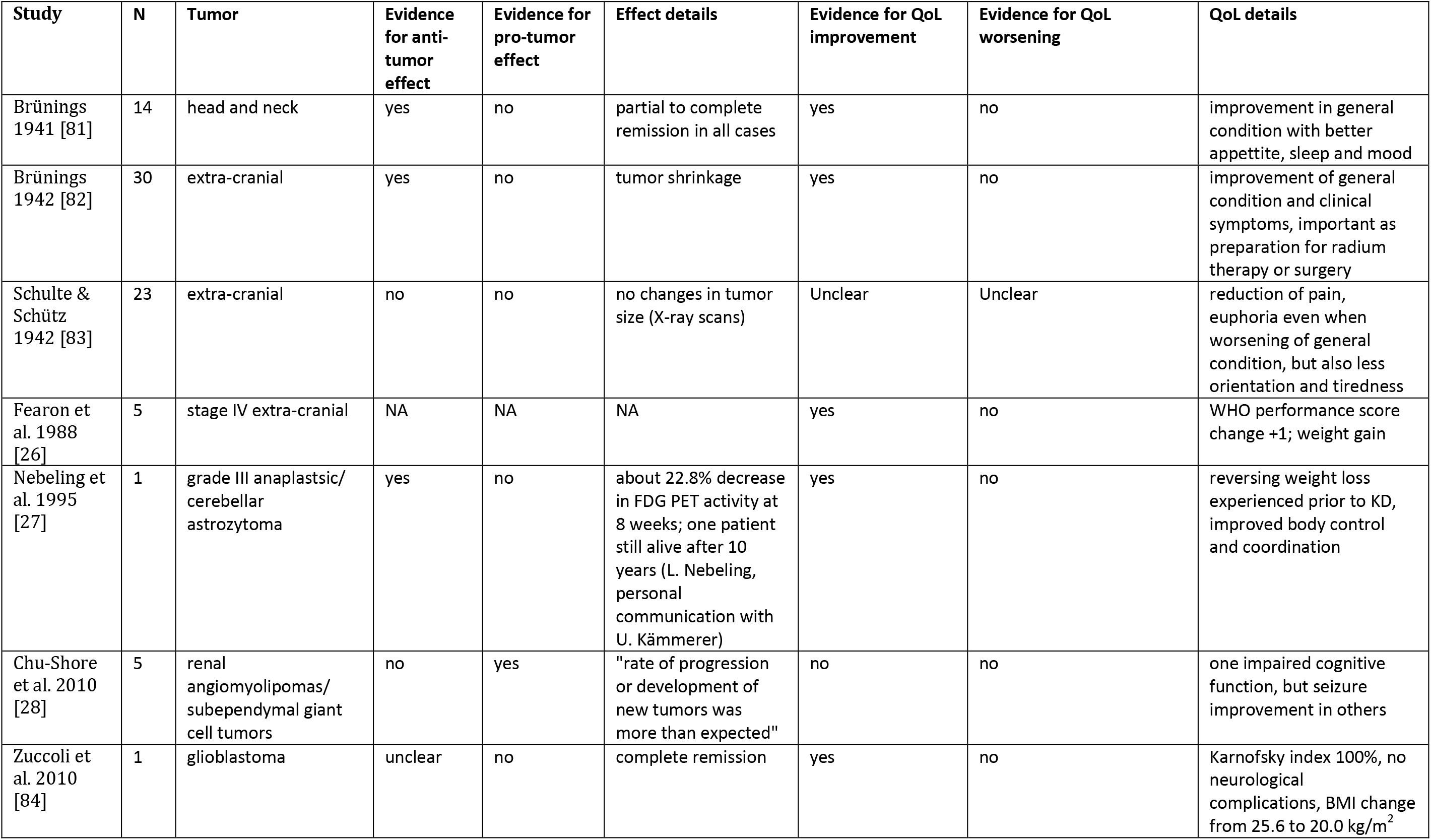
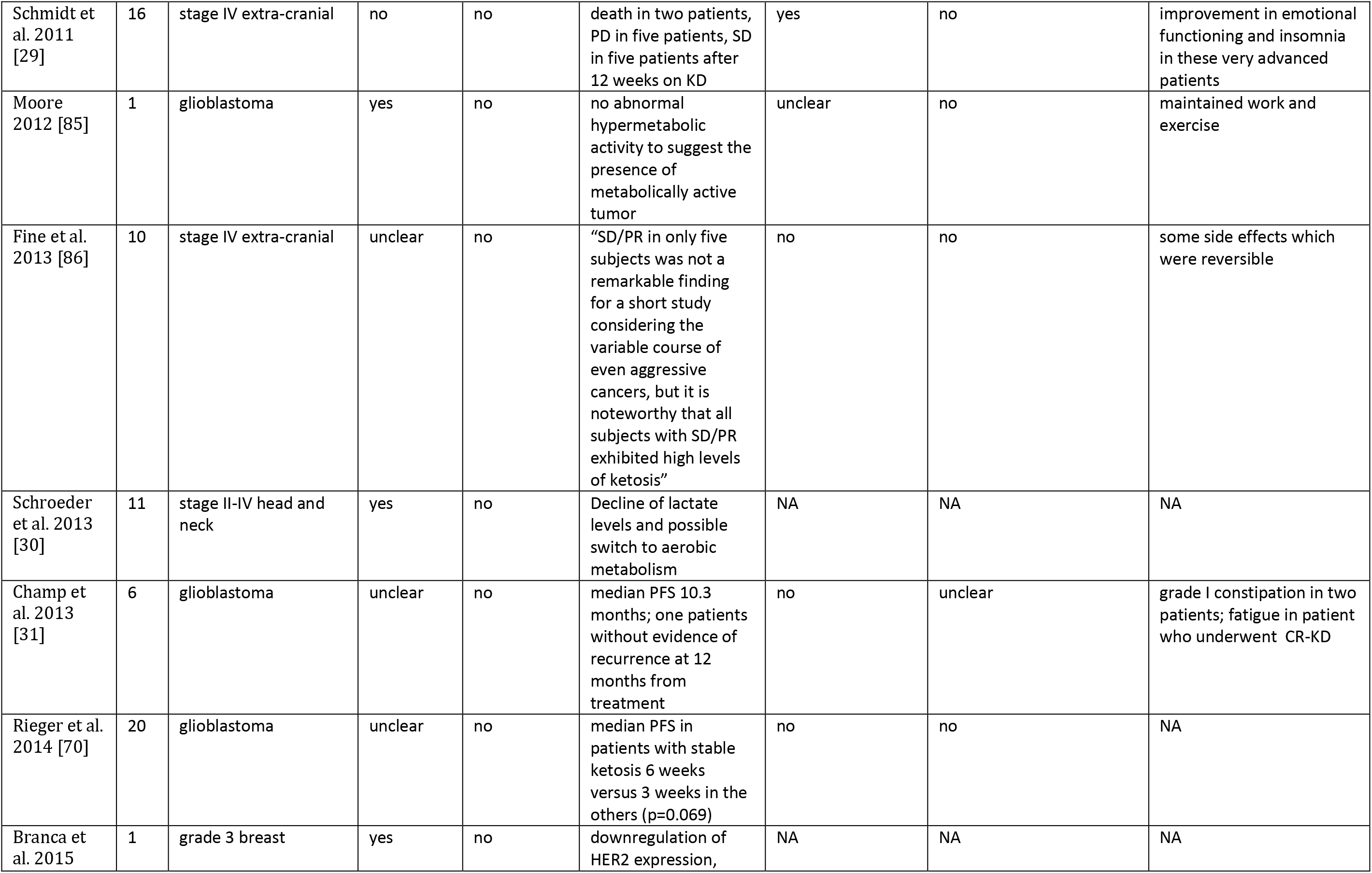
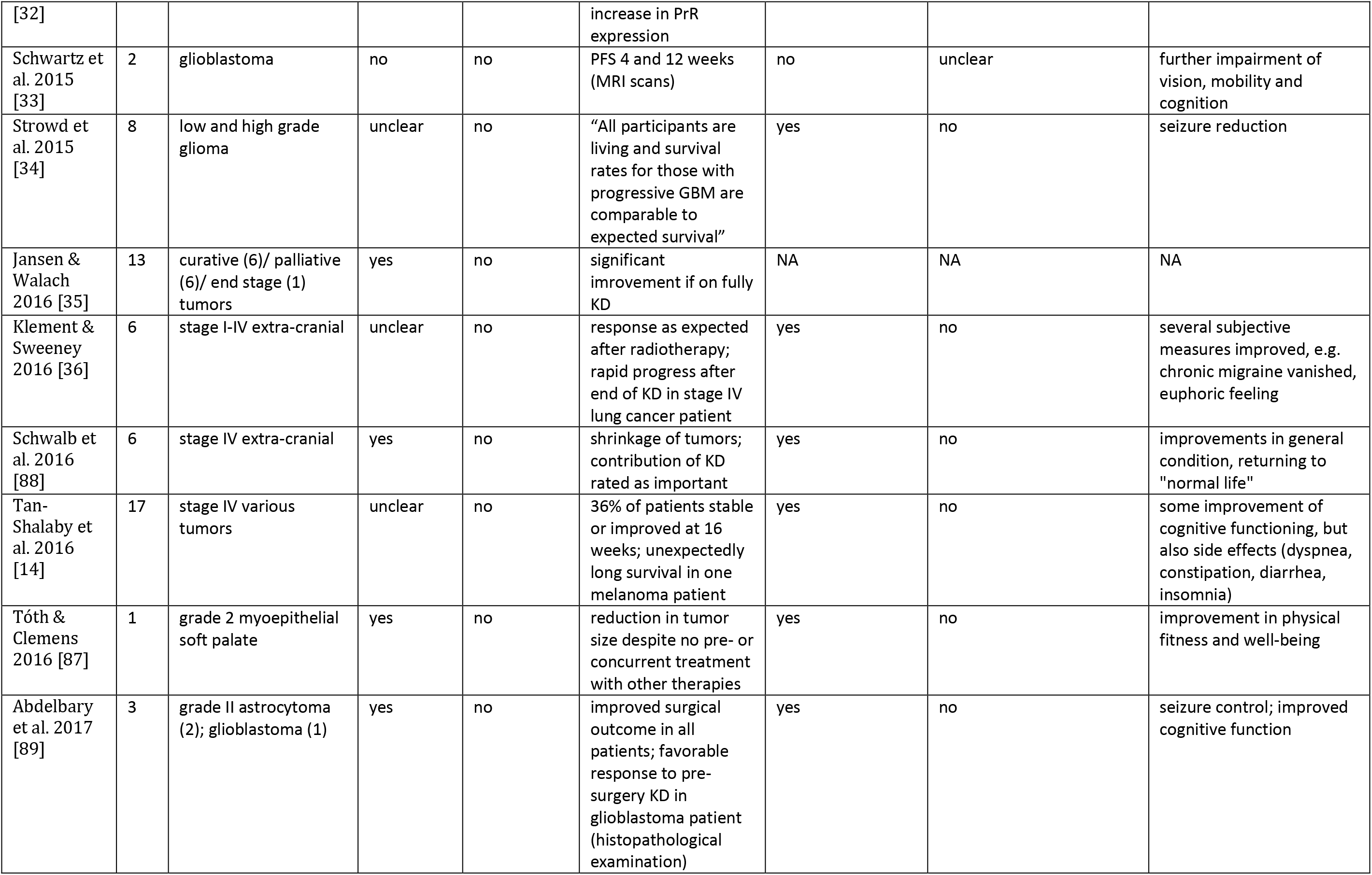
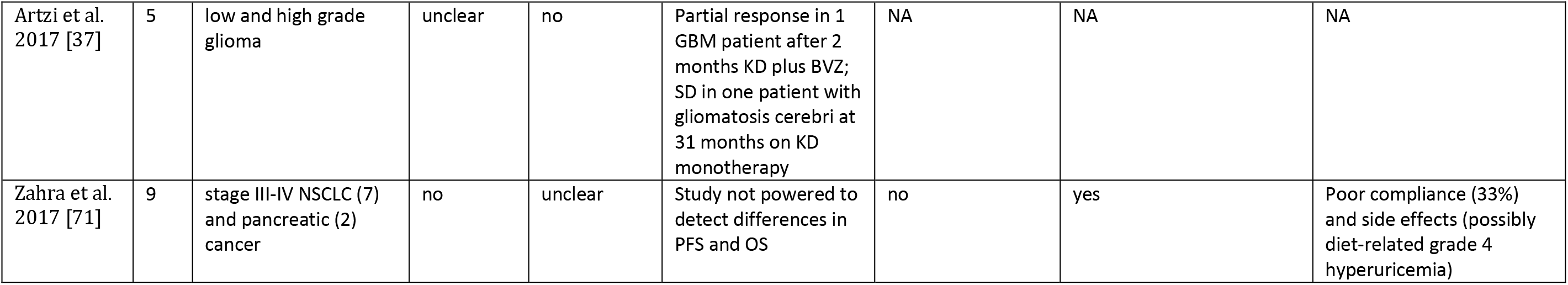
Human studies on KD and cancer. BVZ: Bevacizumab; GBM: glioblastoma multiforme; NSCLC: Non-small cell lung cancer; PD: progressive disease; SD: stable disease

**Table 2.**
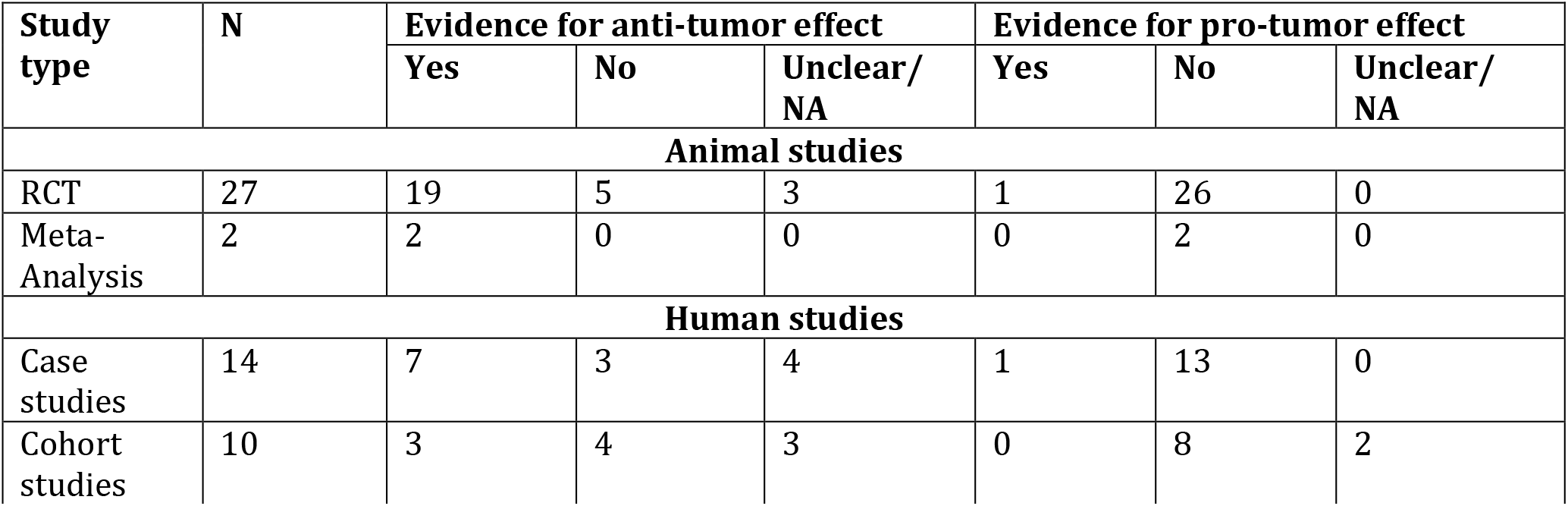
Evidence for the hypotheses that a KD or STS has a positive or negative impact on tumor biology, respectively, each time compared to the hypothesis of no or a neutral impact. “Unclear” evidence is defined as a likelihood ratio close to 1, i.e., neither evidence for the one or the other hypothesis.

| Study type | N | Evidence for anti-tumor effect |  |  | Evidence for pro-tumor effect |  |  |
| --- | --- | --- | --- | --- | --- | --- | --- |
|  |  | Yes | No | Unclear/ NA | Yes | No | Unclear/ NA |
| Animal studies |  |  |  |  |  |  |  |
| RCT | 27 | 19 | 5 | 3 | 1 | 26 | 0 |
| Meta-Analysis | 2 | 2 | 0 | 0 | 0 | 2 | 0 |
| Human studies |  |  |  |  |  |  |  |
| Case studies | 14 | 7 | 3 | 4 | 1 | 13 | 0 |
| Cohort studies | 10 | 3 | 4 | 3 | 0 | 8 | 2 |

In total, 72% (21/29) of the animal studies provided evidence for anti-tumor effects of the KD, either through slower tumor growth or longer overall survival times in the treated animals. Only one study found evidence for pro-tumor effects. Noteworthy, this study [66] provided evidence for both anti-and pro-tumor effects, depending on the length of the KD feeding period. The two meta-analyses concluded that there is overall evidence for anti-tumor effects of KDs in mice [72,73]. Concerning the human data, 42% of the studies (10/24) provided evidence for an anti-tumor effect of KDs. The majority of this evidence came from case reports in which other hypothesis seemed more unlikely as explanations for the observed responses. In seven studies (29%) the evidence was against the KD positively affecting the course of the disease, but only one study revealed evidence for a pro-tumorigenic effect of the KD [28] which is discussed in more detail below.

The results concerning QoL are tabulated in Table 3. Half of the studies (12/24) revealed improvements in QoL, with evidence coming from case and cohort studies with roughly equal contributions. Only one study using an extremely low protein KD with many artificial foods provided evidence for a detrimental effect [71]. Most frequent improvements concerned the general condition [26,27,36,84,85,87,88] and neurological function/seizure control in patients with brain tumors [14,28,34,89]. Side effects attributed to the KD were generally mild and reversible [86]. Weight loss occurred in 11 out of 14 studies in which it was quantified, and was associated with grade II fatigue in one patient who underwent a calorie restricted KD [31].

**Table 3.** Evidence for the hypotheses that a KD or STS has a positive or negative impact on quality of life (QoL), respectively, each time compared to the hypothesis of no or a neutral impact. “Unclear” evidence is defined analogous to Table 2.

| Study type | N | Evidence for positive effect on QoL |  |  | Evidence for negative effect on QoL |  |  |
| --- | --- | --- | --- | --- | --- | --- | --- |
|  |  | Yes | No | Unclear/NA | Yes | No | Unclear/NA |
| Case studies | 14 | 7 | 3 | 4 | 0 | 9 | 5 |
| Cohort studies | 10 | 5 | 3 | 2 | 1 | 7 | 3 |

Three main mechanistic rationales for why researchers chose to implement a KD for cancer patients could be identified (see also [90]):

1. Changes in growth factor and ketone body signaling should affect the complex tumor signaling network, and hence metabolism and growth.
2. The metabolic changes associated with the diet result in alterations of metabolic fuels which are ideally suited for the metabolic demands of the host tissues.
3. KDs increase oxidative stress in tumor cells, making them more vulnerable to oxidative therapies such as radio-and chemotherapy.

### 3.1 Influence on Tumor Cell Metabolism and Growth

#### 3.1.1 Decreasing glucose flux to the tumor

Many tumor cells possess dysfunctional mitochondria and lack certain enzymes necessary for effective ketone body utilization. To compensate, tumor cells would have to rely on substrate fermentation for energy production (reviewed in [2,4]). Many studies mentioned the increased utilization of glucose with subsequent fermentation to lactate by tumors compared to normal tissues, first described by Otto Warburg and co-workers in 1924 [91]. The study authors hypothesized that a KD could be able to limit glucose flux to metabolically inflexible tumors while providing ketone bodies and fat as an alternative fuel for normal tissue [14,27,29–31,33,70,84,85,87,89]. In fact, the very first clinical application of a KD to treat cancer patients was undertaken by the Munich physician Wilhelm Brünings in 1941 based on Warburg's observations and the heuristic that diabetes, hypertensive disease and cancer are all based on an endocrine disturbance of carbohydrate metabolism [81]. Aiming at a drastic reduction of blood glucose levels Brünings developed a “de-glycation method” *(“Entzuckerungsmethode”)* consisting of a diet with less than 50g carbohydrates and three injections of depot insulin per day. In two reports [81,82], he described impressive reductions in tumor size and improvements of QoL that would peak at 2-3 weeks and be sustainable for up to 2-3 months. With more experience, the focus of his treatment apparently shifted towards preparing patients for surgery or radium therapy [82]. While the beneficial effects on QoL could be attributed to the anabolic, analgetic and anti-inflammatory effects of insulin and eventually the high quality of the diet compared to the standard German diet during World War 2, the reductions in tumor growth are harder to understand and could be judged as evidence for an effect specific to the KD. However, subsequently Schulte and Schutz were not able to confirm an anti-tumor effect of Brünings’ method [83].

A KD by itself reduced average blood glucose levels in only a subset of the studies that prescribed no concurrent caloric restriction [26,27,31,86]. Two studies in which patients were advised to always eat to satiety failed to achieve significant reductions in blood glucose levels [36,70], and this was discussed as a possible reason for the failure to halt tumor progression in the study by Rieger et al. [70]. However, more detailed measurements with microdialysis catheters [30] suggest that although the mean plasma concentration remains fairly stable, glucose spikes are greatly reduced under a KD. To also reduce average blood glucose levels, additional calorie restriction was a feasible option in glioblastoma patients [33,84,85,89].

More recent data imply that besides reducing blood glucose levels, a global rise of ketone bodies and free fatty acids is also important for down-regulating glycolysis. Evidence that a KD can reduce tumor glycolysis in some individuals has been gained from studies utilizing FDG-PET scans [14,27,86] as well as microdialysis measurements of tumor lactate concentrations [30]. Jansen and Walach [35] described reductions of the pentose phosphate pathway tumor marker TKTL-1 under a strict KD, providing further evidence for the hypothesis that KDs inhibit tumor glycolysis.

Mechanistically, Fine and colleagues proposed that an “inefficient” Randle cycle [92] occurs in cancer cells upon an inhibition of glycolysis through free fatty acids and ketone bodies; the inefficiency refers to the inability of tumor cells to compensate for the reduced glycolytic ATP production because of mitochondrial dysfunction [92–94]. Palmitic acid, stearic acid and oleic acid were shown to inhibit key glycolytic enzymes in Ehrlich ascites tumor cells [95]. Studies on the anti-tumor effects of oleic acid data back to the 1920s [96,97]; its intake as part of a KD was therefore emphasized by Ruggiero and colleagues, in part also based on the rationale that oleic acid increases the bioavailability of vitamin D_3_[32,88].

Schwartz et al. [33] found that ketolytic enzymes were expressed in tissue specimen from two glioblastoma patients, which indicates that a subset of tumor cells is likely able to metabolize ketones. This was discussed as part of the explanation why treatment with a restricted KD failed in retarding these patient's tumor growth, despite significant reductions in blood glucose and increases in ketone body levels [33].

#### 3.1.2 Changing tumor signaling networks

Several studies emphasized the possibility of a KD to alter several signaling networks in tumor cells [28,34,35,86,88]. One case study described favorable changes in HER2 and progesterone receptor expression during a three week KD supplemented with olive oil and high doses of vitamin D in a breast cancer patient, concluding that both oleic acid and the KD might have worked synergistically to downregulate HER2 expression [32].

The insulin/insulin-like growth factor (IGF) axis has repeatedly been implicated as a major modulator of a complex downstream signaling network in tumor cells including the MAPK and the PI3K-Akt-mTOR pathways [98,99]. Based on this rationale, Fine et al. [86] studied a KD over 4 weeks in ten advanced cancer patients. They found that insulin levels correlated inversely with ketosis, which in turn was correlated to less tumor progression in FDG-PET scans. As the authors themselves stated in their discussion, the outcome *“was not a remarkable finding for a short study considering the variable course of even aggressive cancers”*, so the evidence for the anti-tumor effect hypothesis from this study remains unclear (the likelihood ratio * 1). Cho-Shore and colleagues tested whether KD-mediated mTOR inhibition could slow down growth of tuberous sclerosis complex (TSC) tumors in patients who had started the diet for management of intractable epilepsy [28]. While three of five patients showed improved or complete seizure control, the KD was not able to slow down or inhibit tumor growth; only one patient showed regression of a previously growing subependymal giant cell tumor when he was placed on concomitant treatment with the mTOR inhibitor sirolimus [28]. A subsequent animal model of TSC tumor development showed that while the KD given to Eker rats induced anti-tumor effects in the short term, its long-term (8 months) administration resulted in elevated growth hormone concentrations, MAPK-ERK1/2 and mTOR hyperactivation and accelerated renal tumor growth [66]. This mechanism is consistent with the conclusion of Chu-Shore et al. that in three children with TSC who were on the diet for more than four years, *“the rate of progression or development of new tumors was more than expected in the natural history of TSC”* [28].

### 3.2 Positive influence on body composition

Free fatty acids and ketone bodies are considered to become a major fuel for normal tissues of cancer patients as a consequence of developing insulin resistance [5]. Additionally, ketone bodies have been shown to suppress protein catabolism during starvation [100]. Based on this rationale, Fearon et al. [26] applied a medium chain triglyceride-based KD to five severely cachectic cancer patients for one week. While nitrogen balance was unaltered compared to the previous normal diet, the KD induced a significant weight gain of 2kg and improved their performance status.

In a series of ambulatory patients eating a KD during the course of their radiotherapy in our institution, we found that their relative percentage of fat mass decreased but relative percentage of fat free mass increased while patients lost weight [36]. This was taken as evidence that a KD could positively influence body composition and will be investigated further in a currently running clinical trial [101].

Mechanistically, ketone bodies have been shown to exert anti-cachectic effects by curtailing both the cause and the symptoms of cachexia. In a pancreatic cancer model, the first action was related to the inhibition of glycolytic metabolism in tumor cells, while the second effect was mediated by an *in situ* downregulation of skeletal muscle and adipose tissue degrading proteins [55]. The relation between cachexia and glycolytic tumor metabolism and tumor size is consistent with the classical study by Fearon and colleagues in the Walker 256 sarcoma of the rat in which ketosis was not able to significantly reduce blood glucose levels, tumor growth or weight loss [74]. The significance of the second effect - a direct action of ketone bodies on muscle and adipose tissue - is also consistent with the seminal studies of the MAC16 cachectic mouse adenocarcinoma showing that the maintenance of body weight was greater than anticipated solely from the reduction of tumor size [38].

### 3.3 Synergism with radio-and chemotherapy

Besides impaired oxidative energy production, dysfunctional mitochondria in tumor cells are believed to lead to increased steady-state levels of reactive molecules such as superoxide anion 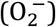 and hydrogen peroxide (H_2_O_2_). Enhanced glycolysis helps to scavenge high steady state levels of these reactive oxygen species (ROS) through increased production of anti-oxidative substrates such as lactate, pyruvate and NADPH [6,90]. A down-regulation of this anti-oxidative defense mechanism, achieved through glucose withdrawal *in vitro* [102] or KDs *in vivo* [51], has been shown to sensitize such tumor cells to additional therapy-induced oxidative stress, while normal cells are left unaffected or even protected [7].

Animal studies provide evidence for the hypothesis that KDs enhance the effects of oxidative stress-inducing therapies such as hyperbaric oxygen [53], radio-and chemotherapy [49,51]. Based on these data, a confirmation of this hypothesis can be found in the case report of a glioblastoma patient showing complete remission after treatment with a restricted KD, radiotherapy and temozolomide [84] as well as our case study of patients undergoing a KD during curative radiotherapy [36] who remain tumor-free after a median follow-up of 112 weeks (unpublished data). However, these data are not evidentially significant because they are not able to distinguish whether the KD contributed to these outcomes or not.

The phase 1 ketolung and ketopan studies were specifically conducted against this background in order to test the feasibility of a 4:1 KD supplying 90% energy from fat and 8% from protein [71]. The diet was highly artificial, with the usual breakfast made from KetoCal^®^ powder and artificial flavorings, as well as low carbohydrate versions of milk, bread and desserts. This KD was poorly tolerated, as only two out of nine patients in the ketolung trial and one of two patients in the ketopan study were able to follow it over the whole course of radiotherapy, and patients experienced several side effects of which at least one, a grade 4 hyperuricemia, was possibly related to the diet. Plasma protein carbonyl content was significantly higher after the KD and radio-chemotherapy intervention, indicating an increase in steady-state levels of ROS-damaged proteins. Again, however, the lack of a control group did not allow any conclusions about the KD contributing to the increased oxidative stress or not.

## Discussion

KDs have been studied in a wide variety of contexts reflecting the broad spectrum of current cancer care. Despite the limitations of these studies which mostly consist of case reports, their data are therefore relevant for judging and comparing hypothesis about the effects of KDs for cancer patients in a real world setting. Earlier reviews have focused on specific mechanisms of the KD [6] or specific contexts in which it is applied [7,9]. This review differs from these works by focusing more broadly on the benefits of KDs for cancer patients within a clear framework of evidence and confirmation.

We can now try to answer the three questions posed in the introduction. The confirmation question was: given the available data-now collated in this review-to which degree should we believe that KDs have beneficial effects for cancer patients? The answer depends on the prior probability that we give to this hypothesis and insofar is always subjective. If the animal studies with their various tumor models evaluated here are taken to estimate a prior probability for the hypothesis that KDs exhibit anti-tumor effects in (at least some) cancer patients, this prior probability appears high (72% of the studies supported anti-tumor effects). Its posterior probability would be even higher, considering that there are human data of some (exceptional) responses to the KD [14,27,89], as well as other data relevant in this context such as the consistent correlations between high blood glucose levels and poor outcome in a variety of human cancers [103–113]. However, a “fundamentalist” sceptic^2^ could always consider such correlations as non-causal (despite mechanistic reasoning suggesting otherwise [114]) and reject the translatability of animal and *in vitro* studies to humans. The published cases with positive responses to the KD would hardly shift such a sceptic's position that there are no anti-tumor effects exerted by the KD. Even in this case, however, the prior probability for the hypothesis that KDs accelerates tumor growth would have to be judged low and the posterior even lower given the lack of human data supporting it. Therefore the hypothesis of an anti-tumor effect of the KD (at least in some patients) is currently confirmed to a much larger degree than the hypothesis of a tumor growth promoting effect. The available data also confirm the hypothesis that KDs are generally safe for cancer patients. In fact, nutritional ketosis is a physiological condition that probably was highly prevalent during human evolution [115,116], providing little *a priori* reason to believe that nutritional ketosis *per se* is unsafe or even dangerous. Again, a fundamentalist sceptic would reject the prior assumption of the safety of a diet appearing extreme compared to dietary guidelines and could point to putative side effects that have been mostly observed in the pediatric population treated with very strict and partially outdated forms of KDs [117].

But is there actual evidence for serious side effects in cancer patients? Serious side effects (grade ≥ 2) reported in the literature are scarce and correlate with extreme versions of the diet, with either concurrent caloric restriction (weight loss [84], fatigue [31]) or extreme macronutrient compositions (hyperuricemia [71]). The ketolung and ketopan studies [71] were important in showing that an artificial 90% fat, 8% protein diet is hardly sustainable. Such a low protein content is at odds with protein requirements of cancer patients which are estimated as at least 1.2g/kg body weight/day with ideal intakes of at least 2g/kg/day for patients in advanced stages [118]. Sufficient intake of high quality protein is important for preserving skeletal muscle mass, the loss of which is much more problematic than that of fat mass [90]. Along these lines, while most of the studies reported weight loss to some extent, this should always be interpreted within the specific study context [117], and for certain patient groups could even be seen as beneficial in terms of QoL and long-term outcome [119] An important finding was that across all settings, half of the studies even provided evidence for an improvement of QoL, an outcome that was rarely mentioned as a primary study goal. Thus, future studies could specifically be designed to investigate which aspects of QoL are influenced by a KD and what constitutes the mechanisms.

Is there evidence for anti-tumor effects in patients? Evidence for the anti-tumor hypothesis is currently limited and of low strength in the human data (Table 2). This is partly based on the design of the studies undertaken so far that does not allow discriminating between the antitumor hypothesis and the hypothesis of a neutral or no effect. From a realist standpoint, the biochemical events that determine the effects of a KD on tumor cells under different contexts or boundary conditions are *ontological aspects* that are essential for explaining and predicting individual patient outcomes [20]. Unfortunately the causation question concerning a mechanistic explanation of observed study outcomes has been left unanswered in most cases. While most researcher agree on the causal importance of lowering blood glucose levels and increasing ketone body levels for achieving a therapeutic effect, there are still too many unknowns in the inferential chain linking these metabolic changes to patient-relevant outcomes, and currently running controlled trials will be of great value for filling these knowledge gaps [9].

Finally, given the above answers to the confirmation and evidence questions, which advice can be given to cancer patients and their physicians (the *decision* question)? It is clear that with many cancer patients wishing to try a ketogenic diet and seeking professional guidance to do so safely, we cannot wait until results of RCTs have been obtained (if they ever will) [120]. Patients should be informed about the totality of evidence including plausible mechanistic reasoning but the current limitations of evidence from human data, a situation typical for an emerging new treatment. The data implicate that a logical argument can be derived stating that KDs could safely be offered to patients, since the probability of potential benefits appears much larger than the probability of serious side effects. Of course this includes the assumption of a setting in which patients are both intrinsically motivated and externally supervised, and the diet fulfills the protein and micronutrient requirements. Again, this judgement is largely subjective and more objective evidence needs to be gathered from further mechanistic studies, larger case series and – most importantly – controlled clinical trials.

A limitation of this review is that one could question the assignment of evidence for the antitumor effect hypothesis to particular study data as given in Table 1. I tried to objectively evaluate alternative hypothesis appearing realistic within the context of each particular study, but did not consider any form of *ad hoc* hypothesis such as a spontaneous remission as likely. The comments on effect details in Table 1 have been included to justify my assignment of evidence. The definition of evidence that I used could also be criticized. For example, Cartwright et al. find *“probabilistic relations between the evidence and the hypothesis…not useful for evidence-based policy”* [121], while at the same admitting that we have no practicable theory of evidence. Again, their statement exemplifies the confusion between evidence and data. The distinction between evidence, confirmation and data used herein has been shown to overcome a variety of scientific and philosophical paradoxes [15] and distinguishes this work from others that refer to evidence without having a clear concept of this term. The evidential conclusions drawn from the studies are robust against other measures of evidence which incorporate the comparison between two hypotheses.

As a cautionary note, some recent animal experiments have revealed tumor growth stimulation through administration of ketone bodies [68,122,123]. Two studies using breast cancer models showed that BHB infusions accelerated tumor growth by serving as an energetic substrate for oxidative (non-Warburg) tumor cells [122,123]. Since no KD was applied, no evidence for or against its use can be derived from these data. The study of Xia et al. [68] showed that acetoacetate, but not BHB, accelerated tumor growth of BRAF V600E-expressing melanoma xenografts, which led the authors to express concerns about KDs for patients with tumors that harbor such mutations. However, in this study a KD did only increase acetoacetate, but not BHB levels, which makes this KD appear different from all other animal experiments and questionable as a model system for a KD applied to humans where ketosis is characterized by more than four-fold lower acetoacetate than BHB levels [124]. It is also noteworthy that the best response to a KD in the study by Tan-Shalaby et al. [14] was seen in a patient with BRAF-V600 positive stage V melanoma who stayed tumor-free after surgery and a prolonged KD at 131 weeks of follow-up. Based on these animal data, the possibility that a subset of human tumors could be stimulated by a KD should therefore be considered, but is not justified by the human data published so far and not even supported by evidence from these studies themselves (because a KD was not used in two of them and elicited completely different effects to a KD in humans in the third).

## CONCLUSIONS

KDs have been studied in humans in a wide variety of contexts that are representative of modern cancer treatment. Due to a lack of controlled trials the human data are only able to provide very low level evidence for anti-tumor effects of KDs which is also limited to individual cases. The total evidence is upgraded to some extent by mechanistic reasoning. Nevertheless, some exceptional responses have justified the belief that KDs can exert antitumor effects at least for some patients because the posterior probability for this hypothesis is higher than for its alternatives. Furthermore, there is no evidence or reason to believe that KDs have serious side effects or would accelerate tumor growth. The logical conclusion is therefore that KDs are promising and worth of further study since the probability of achieving an anti-tumor effect is apparently greater than that of causing serious side effects.

## COMPLIANCE WITH ETHICAL STANDARDS

### Ethical approval

This review was undertaken without requesting ethical approval, since ethical approval was granted for most of the included studies, all human data were anonymous, and the ethical recommendations of the Helsinki Declaration were adhered to.

## Footnotes

1 As most philosophers of science, Carnap [16] who formulated this principle made no clear distinction between evidence, data and confirmation [15]. Within our framework (see Materials and Methods section), his principle of total evidence would better be described as the principle to consider all relevant available data when making inferences about hypotheses.

2 A distinction should be made between “fundamentalist” skepticism, a term coined by M.M. ℆irkoviℇ [125] and other, more rational forms of skepticism that for example acknowledge the mechanistic data, but consider the human data not supportive for methodological reasons. Examples of the former have been critizised by us previously [117,126].

